# scShaper: ensemble method for fast and accurate linear trajectory inference from single-cell RNA-seq data

**DOI:** 10.1101/2021.05.03.442435

**Authors:** Johannes Smolander, Sini Junttila, Mikko S. Venäläinen, Laura L. Elo

## Abstract

Computational models are needed to infer a representation of the cells, i.e. a trajectory, from single-cell RNA-sequencing data that model cell differentiation during a dynamic process. Although many trajectory inference methods exist, their performance varies greatly depending on the dataset and hence there is a need to establish more accurate, better generalizable methods. We introduce scShaper, a new trajectory inference method that enables accurate linear trajectory inference. The ensemble approach of scShaper generates a continuous smooth pseudotime based on a set of discrete pseudotimes. We demonstrate that scShaper is able to infer accurate trajectories for a variety of nonlinear mathematical trajectories, including many for which the commonly used principal curves method fails. A comprehensive benchmarking with state-of-the-art methods revealed that scShaper achieved superior accuracy of the cell ordering and, in particular, the differentially expressed genes. Moreover, scShaper is a fast method with few hyperparameters, making it a promising alternative to the principal curves method for linear pseudotemporal ordering. scShaper is available as an R package at https://github.com/elolab/scshaper.

## 1 Introduction

Single-cell RNA-sequencing (scRNA-seq) is a powerful technology for studying dynamical processes of cells in tissues (Tanay and Regev, 2017). Computational models, known as trajectory inference methods, are needed to infer a representation of the cells, i.e. a trajectory that models the real phases at which the cells are developing during the process. The workflow of trajectory inference can be roughly divided into three main steps: 1) preprocessing that includes quality control to remove lowly expressed genes and poor quality cells, normalization and dimensionality reduction; 2) topology inference that typically involves clustering and building a graph based on the clustering, such as a minimum spanning tree (MST); 3) generation of a pseudotime for each lineage of the trajectory, i.e. a set of continuous values ranging from 0 to 1 that measures the progression of the cells along the lineage. While many methods have been developed for trajectory inference (Saelens et al., 2019), recent works have mainly attempted to improve the topology inference (Todorov et al., 2020; Cao et al., 2019). As topology inference is very challenging, users often need to try several different tools to find a suitable topology. However, in many cases a simple linear model can be sufficient, and therefore more accurate linear trajectory inference methods are still needed as well.

Linear trajectory inference is strictly limited to the linear topology that does not allow branching or cycles. For estimating the pseudotime, the most common method currently is the principal curves method, and it was recently shown that the current best linear trajectory inference methods, SCORPIUS and Embeddr (Cannoodt et al., 2016; Campbell et al., 2015), use the principal curves method (Saelens et al., 2019). Slingshot (Street et al., 2018), which was ranked as the best tree-based method in the comparison study, also utilizes the principal curves method. The principal curve is a smooth one-dimensional curve that passes through the middle of a p-dimensional dataset (Hastie and Stuetzle, 1989). The algorithm starts from a prior curve, which is by default set to the first principal component, and then proceeds to iteratively project the samples using a smoothing function so that the samples pass through the middle of the dataset. However, it is known that the regular principal curves algorithm can perform poorly with more complex trajectories, such as spiral trajectories (Ozertem and Erdogmus, 2011), and for this reason some scRNA-seq trajectory inference analysis methods have attempted to address this limitation using other initializers than the first principal component. SCORPIUS (Cannoodt et al., 2016), which was ranked as the best linear trajectory inference method in the recent comparison study (Saelens et al., 2019), begins by first clustering the dataset using the k-means algorithm and then infers the shortest path through the clustering and uses this as prior in the principal curves method. However, this approach raises several issues, such as how to select the number of clusters k and the sensitivity of the shortest path to the clustering. Moreover, it is not guaranteed that the subsequent iterative smoothing converges to a final curve that correlates well with the shortest path.

To address these limitations, we introduce scShaper, a new trajectory inference method that enables accurate linear trajectory inference from scRNA-seq data using an ensemble approach that combines multiple pseudotime solutions into a single more accurate ensemble solution. scShaper is based on graph theory and solves the shortest Hamiltonian path of a clustering, utilizing a greedy algorithm to permute clusterings computed using the k-means method to obtain a set of discrete pseudotimes. In contrast to other methods that rely on a single clustering, scShaper clusters the dataset multiple times using a range of different k values and combines the resulting discrete pseudotimes into a solution, which is less sensitive to the number of clusters. scShaper also excludes trajectories that are considered outliers and builds the solution based on the largest subset of mutually correlating pseudotimes, effectively mitigating the instability of the single clustering -based approach.

To demonstrate the benefits of scShaper, we repeated the same benchmarking as in the recent study, which compared 60 different scRNA-seq trajectory inference tools in a comprehensive manner (Saelens et al., 2019). The results suggested that scShaper was the best method in terms of the overall performance that measures accuracies of cell ordering and differentially expressed features. In particular, there was a considerable improvement in the accuracy of the differentially expressed features, suggesting that scShaper is able to infer trajectories that more accurately account for different cell subpopulations. Moreover, we compared scShaper with the regular principal curves method under the same preprocessing steps using the same benchmarking and found that the performance of scShaper was again statistically significantly better, suggesting that the performance boost is not only attributable to the different preprocessing steps of the trajectory inference methods, but the differences in the pseudotime inference. Conveniently, the run time of scShaper was also comparable with the state-of-the-art methods. Finally, we demonstrate that scShaper is also able to infer accurate trajectories for a set of different nonlinear trigonometric trajectories, most of which were too complex for the regular principal curves algorithm, suggesting scShaper could become a well-generalizable alternative to the principal curves method for inferring linear paths through datasets.

## 2 Methods

### 2.1 Linear trajectory inference algorithm of scShaper

In the following, we describe the four steps of the linear trajectory inference algorithm of scShaper that starts with preprocessed data.

#### 1. Discrete pseudotime estimation using shortest Hamiltonian path permuted clustering

As the basis of scShaper, we first describe an unsupervised clustering and graph theory -based algorithm that can be used to relabel clustering so that the numeric cluster labels correlate with the path through the middle of the data (**Figs. 1a-d**). After clustering, the labels are initially random without any dependency between the labels and neighbourhoods of the clusters (**Fig. 1a**). Here, we refer to such a relabeled clustering that correlates with the path as the discrete pseudotime (**Fig. 1d**). To generate a discrete pseudotime, we need to find the permutation of the clustering that minimizes the distances between the cluster centroids of the adjacent clusters (**Figs. 1b-c**). From a mathematical standpoint, this is a graph theory problem, in which the clusters are the vertices and the connections between the clusters are the edges weighted by the distances between the clusters, and we seek the shortest path that visits each vertex exactly once. This kind of path, in which each vertex is visited only once, is called a Hamiltonian path, and the problem can be described as finding the shortest Hamiltonian path. However, finding the shortest Hamiltonian path is known to be NP-hard, and hence a greedy algorithm is necessary to achieve a high computational performance with large numbers of clusters. In the following, we describe a pseudocode (**Algorithm 1**) for our greedy algorithm, which is essentially Kruskal’s algorithm for finding the minimum spanning tree (MST) of a graph. The difference is that a Hamiltonian path has slightly stricter limitations than MST, since the maximum degree of a vertex, i.e. the maximum number of adjacent edges, is two. Therefore, the problem can also be characterized as a degree-constrained MST problem. As in Kruskal’s algorithm, the greedy algorithm adds edges into the graph by starting from the shortest distance and discards connections that form cycles, but also connections that exceed the degree constraint.

**Figure 1.**
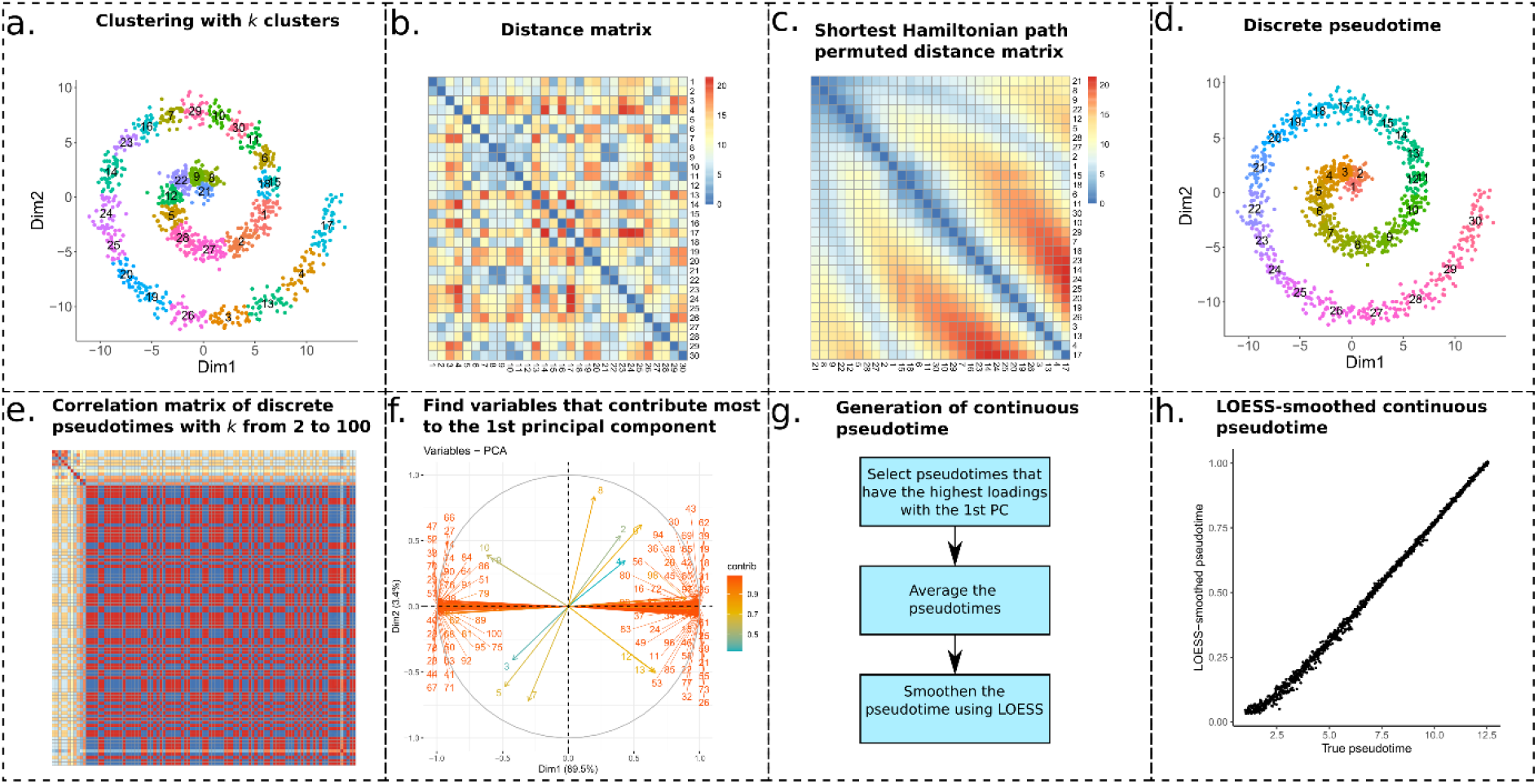
Schematic of the linear trajectory inference method of scShaper. **(a)** The data points are clustered using the k-means algorithm into k clusters. **(b)** A distance matrix is calculated between the cluster centroids. **(c)** Sort the clusters by solving the shortest Hamiltonian path problem using a greedy algorithm. **(d)** To create a discrete pseudotime, rename the original cluster labels to the permutation which rearranges the optimal permutation into ascending or descending order. **(e)** Repeat steps **a-d** for k values from 2 to 100 to obtain a set of discrete pseudotimes. The subplot visualizes a correlation matrix of the discrete pseudotimes. Red, yellow and blue colors denote high positive, low and high negative correlation, respectively. **(f)** Perform principal component analysis (PCA) to determine the largest subset of linearly correlating pseudotimes. **(g)** Schematic of the process that generates a continuous pseudotime from the discrete pseudotimes, which includes finding the pseudotimes that have the highest loadings with respect to the first principal component, averaging the selected pseudotimes and finally applying local regression (LOESS) to smoothen the pseudotime. **(h)** Comparison of the ground truth and estimated continuous pseudotimes for the spiral trajectory.

##### Algorithm 1: A greedy algorithm for finding a solution to the shortest Hamiltonian path problem.

1. **Input:** A connected weighted graph *G* = (*V_G_*, *E_G_*) with *k* vertices and the edge weights *D* = (*d_ij_*), where *d_ij_* is the Euclidean distance between the centroids of clusters *i* and *j*.
2. **Output:** A graph *S* = (*V_G_, E_S_*) that provides a solution to the shortest Hamiltonian path.
3. **Initialization:** Sort the edges of *G* = (*V_G_, E_G_*) by their weights into ascending order. The total number of edges in G is *N* = (*k*^2^ – *k*)/2 after removing inverse edges and self-connections. Let *e*_1_ be the edge with the smallest weight and set *E_S_* = {*e*_1_}.
4. **for** (*i* in **range**(2,*N*)):
5. **if** adding edge *e_i_* to graph *S* = (*V_G_, E_S_*) keeps *degree*(*S*) ≤ 2  **and** forms no cycles to *S*:
6. **add** *e_i_* to *S*
7. **if** |*E_S_*| = *k* – 1:
8. **return** *S*

After finding a solution to the shortest Hamiltonian path problem, we obtain a permutation that finds the shortest path through the cluster centroids (**Fig. 1c**). Next, the discrete pseudotime (**Fig. 1d**) is obtained by renaming the original cluster labels (**Fig. 1a**) to the permutation that rearranges the optimal permutation into ascending or descending order.

Although the approach can be applied to any clustering algorithm to find a discrete pseudotime, we chose k-means due to its high computational efficiency and algorithmic simplicity. In our scShaper R package, we use the k-means implementation from the stats R package.

#### 2. Estimate discrete pseudotime for a range of k values

A single discrete pseudotime that is constrained to a single number of clusters (k) is not particularly useful for scRNA-seq trajectory inference. Instead, we would like to have a continuous pseudotime that correlates with the path through the middle of the input data. Additional issues that might arise from using a single pseudotime are that the solution can be sensitive to the stochasticity of the k-means clustering and there is no guarantee that the absolute shortest path is found using the greedy algorithm, which can consequently lead to a suboptimal discrete pseudotime. To address these limitations, we estimate the discrete pseudotime using a range of different k values (**Fig. 1e**), *k* ∈ [*k_min_*, …, *k_max_*], where *k_min_* and *k_max_* are the lower and upper limits of the number of clusters k, respectively. An ensemble solution is created based on these clusterings, resulting in a continuous and more robust solution than a single discrete solution. Here, we always use *k_min_* = 2 and shall constrain *k_max_* in **Section 3.1**.

#### 3. Principal components analysis

After finding a set of discrete pseudotimes for a range of different k values, scShaper performs PCA to analyse linear dependencies between the pseudotimes (**Figs. 1f-g**). We calculate the first two principal components, with feature standardization, and determine the pseudotimes that contribute most to the first principal component based on the feature loadings, which are subsequently used as the basis for generating the final continuous pseudotime. The next step is to average the pseudotimes to generate a crude initial pseudotime. The discrete pseudotimes are min-max scaled and ordered in the same direction using the PCA loading information, after which the pseudotime values for each cell are averaged. To perform PCA, we use the prcomp function from the stats R package.

#### 4. Local regression smoothing

As the final step, we apply local regression (LOESS) using the stats R package to smoothen the crude average pseudotime, where the average pseudotime and its ranking are the response and predictor variables of the model, respectively. LOESS has two important parameters: “span” that controls the degree of the smoothing (default 0.75) and “degree” that determines the degree of the polynomials (default 2). We left the polynomial degree to its default, but investigated how the span parameter affected the performance in **Section 3.1**.

### 2.2 Preprocessing before trajectory inference

As with all scRNA-seq trajectory inference methods, several preprocessing steps are required before the actual trajectory inference with scShaper, including cell normalization and dimensionality reduction by feature selection, which are typically left for the user to decide. Since the benchmarking datasets (**Section 2.5**) were already partially preprocessed by the authors of the comparison study, including filtering outlier cells, normalization and feature selection, the only remaining step in this study was the dimensionality reduction. PCA followed by t-distributed stochastic neighbour embedding (t-SNE) (Maaten and Hinton, 2008) is perhaps the most common dimensionality reduction workflow in scRNA-seq data analysis, and we therefore selected that as the default dimensionality reduction method. To perform combined PCA and t-SNE analysis, we used the Rtsne package with default parameters, including 50 principal components and the perplexity value of 30, with the exception of a three-dimensional instead of a two-dimensional output.

### 2.3 Additional steps after trajectory inference

After trajectory inference, the trajectory is usually visualized as a straight line through a two-dimensional visualization generated by a non-linear dimensionality reduction method, such as t-SNE, UMAP or multidimensional scaling (MDS). The user is typically also interested in investigating differential expression along the trajectory. Conveniently, the dyno R package (Saelens et al., 2019) offers all these functions and supports integration with any trajectory inference method. We provide the scShaper R package together with a vignette, which integrates scShaper with commonly performed preprocessing steps (**Section 2.2**) and dyno to perform a full-scale trajectory inference analysis.

### 2.4 Simulation of trigonometric trajectories

We simulated several trigonometric trajectories, which are summarized in **Supplementary Table 1**, to constrain the parameters of scShaper (**Section 3.1**), including 2D spirals, quadratically growing 2D spirals, 3D spirals, quadratically growing 3D spirals, and sine waves. Using the trigonometric trajectory functions and a sequence of t values with increments of 0.01, we, generated datasets with 1157 to 1785 samples. In addition, we simulated several trajectories with each function by adding a varying amount of Gaussian noise into the variables by adjusting the standard deviation of the Gaussian distribution from 0.05 to 0.95. After adding the noise, the exact ground truth pseudotime becomes unknown, but the t variable still provides a good approximation of the ground truth. The analysis of these trajectories provides a means to constrain the parameters of scShaper in a manner that was not biased towards the scRNA-seq benchmarking.

### 2.5 Benchmarking

To benchmark scShaper for trajectory inference of scRNA-seq data against current state-of-the-art methods, we used a framework from a recent comparison study (Saelens et al., 2019), which benchmarked over 60 different methods using a wide array of different datasets, both simulated and real, and 30 different performance metrics.

The benchmarking data included in total 69 linear datasets, of which 39 and 30 were real and simulated datasets, respectively. The real datasets included gold and silver datasets. For the gold datasets, the discrete pseudotime is known, i.e. the different cell types and the order in which they differentiate. The silver standard datasets include a continuous pseudotime, which were inferred using scRNA-seq trajectory inference methods by the authors of the original studies. The simulated datasets were simulated using four different simulators: dyntoy (Saelens et al., 2019), dyngen (Saelens et al., 2019), PROSSTT (Papadopoulos et al., 2019) and Splatter (Zappia et al., 2017), and provided the most accurate ground truth.

Of the twelve linear trajectory inference methods that were benchmarked in the comparison study (Saelens et al., 2019), we selected four methods for our comparisons. SCORPIUS (scorpius) (Cannoodt et al., 2016), Component 1 (comp1) and Embeddr (embeddr) (Campbell et al., 2015) were the three most highly ranked methods, whereas the fourth selected method, ElPiGraph Linear (elpilinear) (Albergante et al., 2020), is the only method that uses some form of ensemble learning, although much different compared with the approach of scShaper. SCORPIUS and Embeddr use the principal curves method in the actual pseudotemporal ordering step, but they differ in their approach to dimensionality reduction: SCORPIUS uses MDS, whereas Embeddr uses Laplacian eigenmaps. Component 1 is the simplest algorithm, which only computes the first principal component and uses it directly as the pseudotime. ElPiGraph Linear is an extension of ElPiGraph, which is limited to linear trajectories. ElPiGraph has its own considerably more unique approach based on elastic principal graphs, which resembles the principal curves method, but also allows branching to model more complex topologies. ElPiGraph also utilizes ensemble learning to build a more accurate consensus solution based on multiple principal graphs, making it hence a good method to compare with scShaper.

To measure the overall performance of the methods, we followed the dynbenchmark benchmarking framework of the previous comparison study, which used the geometric mean of four metrics that measure 1) the accuracy of cell ordering (cordist), 2) the accuracy of the differentially expressed features (wcor), 3) the accuracy of the topology inference (HIM), and 4) the accuracy of the branching points (F1_branches), of which the last two are always constant with value 1 (perfect) in linear trajectory inference. Therefore, in linear trajectory inference the overall performance was determined by cor_dist and wcor so that low scores in either metric were penalized by the geometric mean. Here, cordist quantifies the similarity between the known and predicted trajectories in terms of correlation of pairwise distances between the two trajectories, and wcor quantifies the agreement between the differentially expressed features obtained using the known and the predicted trajectories. More detailed descriptions of the measures are given in the documentation of dynbenchmark (https://github.com/dynverse/dynbenchmark).

To determine whether the observed performance boost for scShaper was statistically significant, we used the Wilcoxon signed-rank test from the stats R package. The related benchmarking code is available online at https://github.com/elolab/scShaper-benchmarking.

### 2.6 Software and hardware for measuring run time

The run times of the five benchmarked methods were measured on a laptop with Ubuntu 16.04 LTS operating system, 2-core 2.30 GHz Intel(R) Core (TM) i5-6200U processor and 8 GB DDR4 of RAM. As data we used three datasets with 1000, 5000 and 10000 cells, each with 1000 features, simulated using the dyntoy tool.

## 3 Results

### 3.1 Constraining parameters of scShaper

To constrain the parameters of scShaper, we considered 50 different datasets (**Section 2.4**) that were simulated from five different trigonometric functions with varying levels of Gaussian noise added into the variables. We ran scShaper with different parameter configurations and calculated the root mean squared error (RMSE) and the Pearson correlation coefficient between the approximative ground truth and estimated pseudotime for each dataset.

The results suggest that the two *k*-means clustering related parameters, the maximum number of clusters (**Figs. 2a,d**) and the number of random sets in *k*-means (**Figs. 2b,e**) had significant effects on the performance of scShaper. scShaper performed better with a higher maximum *k* value, which is intuitive considering this adds more information into the ensemble set. However, as increasing the upper limit of the *k*-value also increases the run time (**Supplementary Fig. 1**), we concluded that *k* = *100* would be a good trade-off between the run time and the performance. The number of *k*-means initializations had an inverse effect on the performance, where one initialization outperformed hundred initializations. This occurs due to the fact that increasing the number of initializations increases the robustness of the clustering. However, our algorithm benefits from less robust clustering, because this way we obtain more dissimilar discrete pseudotimes, which subsequently provides more information about the relative positions of the cells in the trajectory. Since using a smaller value also helps to minimize the run time of scShaper, we set the default value of the number of *k*-means initializations to 1.

**Figure 2.**
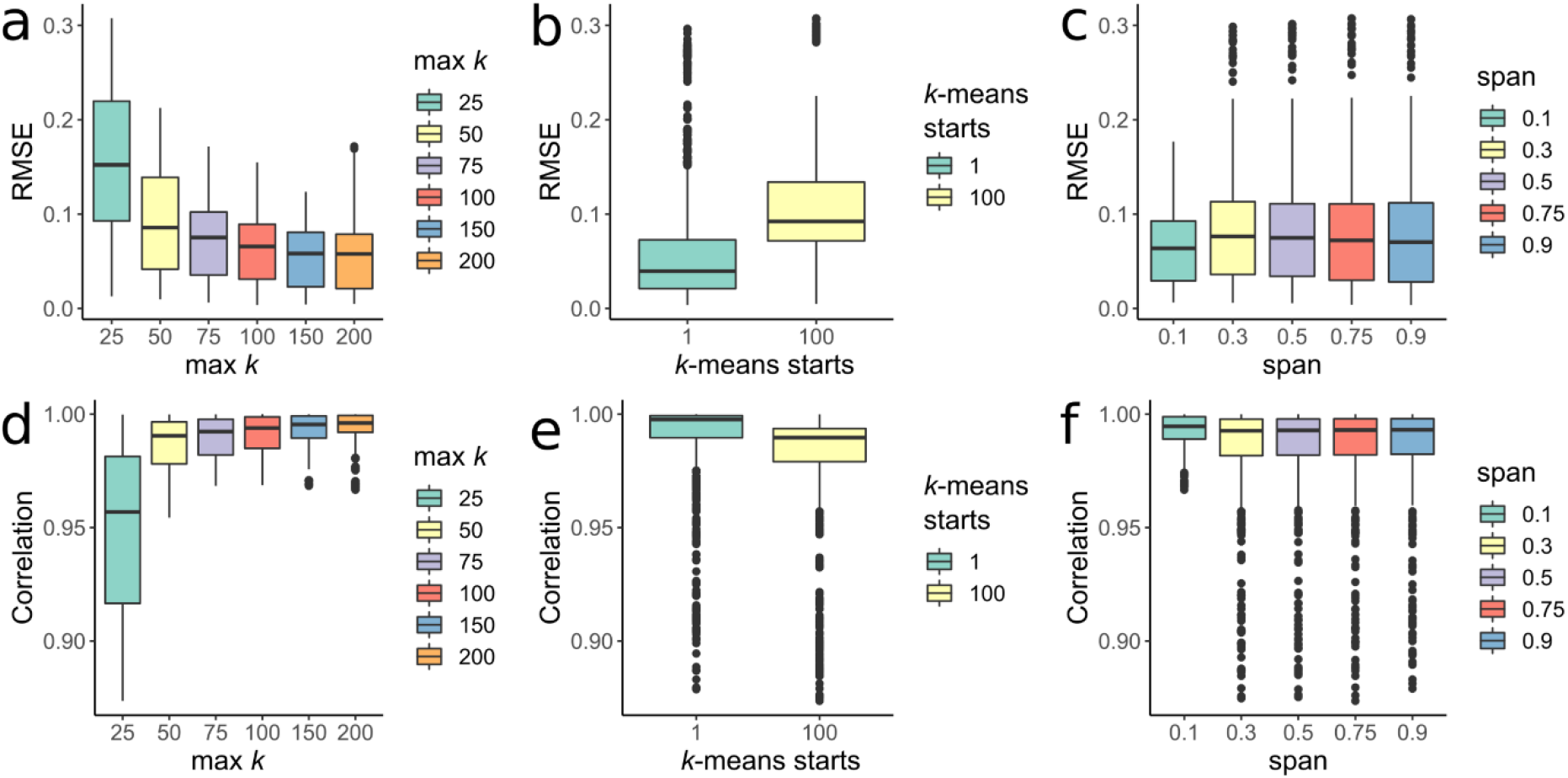
Analysis of simulated trigonometric trajectories to constrain the three hyperparameters of scShaper. **(a,d)** The maximum number of clusters for which the discrete pseudotime was calculated using the k-means algorithm, the minimum being 2. **(b,e)** The number of k-means initializations. **(c,f)** The span parameter that controls the degree of smoothing in Local Polynomial Regression Fitting (LOESS). y-axis in the subplots **a-c** and **d-f** signify the root mean squared error (RMSE) and the Pearson correlation coefficient, respectively, calculated for all the datasets **(Section 2.4**) between the ground truth and inferred pseudotimes.

The third and final parameter, span, controls the degree of smoothing in LOESS, where a higher value smoothens the curve more. Unlike with the two *k*-means related parameters, the differences between the performances obtained using different span values were relatively small (**Fig. 2c,f**). Because the analysis suggested that a lower degree of smoothing was, in general, better, we set the default value of span to 0.1.

### 3.2 Benchmarking using scRNA-seq data

We benchmarked scShaper against four state-of-the-art scRNA-seq trajectory inference methods, SCORPIUS (scorpius), Component 1 (comp1), Embeddr (embeddr) and ElPiGraph Linear (elpilinear), using the current best practices (**Section 2.5**). **Figure 3** visualizes the main results of the benchmarking. The results suggested that the cell ordering (cordist) of scShaper was at least as accurate as of the four other methods (**Fig. 3a**; *p* < 0.05 compared to comp1, elpilinear and scorpius; *p* = 0.1 compared to embeddr). The second correlation metric (wcor), measuring the accuracy of the differentially expressed features, was consistently higher for scShaper than for the other methods (**Fig. 3b**; *p* < 0.01 compared to all other methods). Similarly as in the original comparison study, we calculated the geometric mean of these two metrics to determine the overall score (**Fig. 3c**). The results suggested again that scShaper was the best method (*p* < 0.05 compared to all methods).

**Figure 3.**
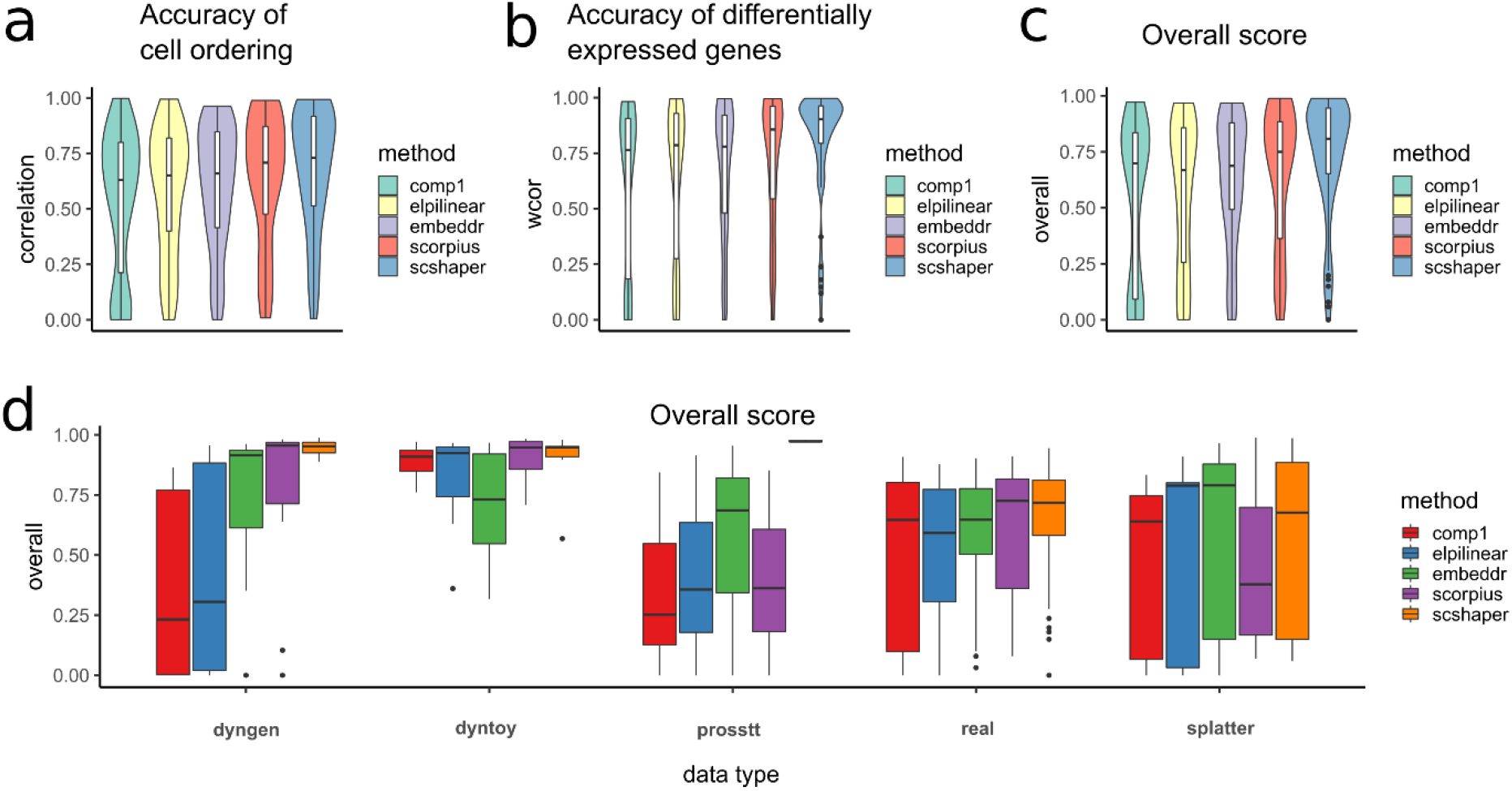
Benchmarking results for scRNA-seq data. **(a)** Accuracy of the cell ordering (cordist). **(b)** Accuracy of the differentially expressed features (wcor). **(c)** The overall score based on the geometric mean of the two previous metrics. **(d)** The overall scores grouped by data type, where dyngen, dyntoy, prosstt and splatter are simulators and real denotes real data.

Next, we grouped the overall scores by the type of the data, i.e. which simulator was used or if the data were real. According to these results (**Fig. 3d**), scShaper showed superior performance for three of the simulators (dyngen, dyntoy, prosstt) and also yielded the highest average performance for the real datasets. For dyngen, dyntoy and real datasets the margins between the medians of SCORPIUS and scShaper were small, but compared to SCORPIUS, scShaper had noticeably fewer datasets with low performance. The splatter datasets were a clear exception in the sense that there were several datasets, which none of the methods were able to model accurately. The performance of scShaper for splatter was moderate, whereas SCORPIUS exhibited inferior performance.

The whole comparison using the dynbenchmark framework included in total 31 benchmarking metrics, the remaining of which are visualized in **Supplementary Figures 2-4**. Most of these metrics show similar rankings for the methods, where scShaper is consistently among the best methods.

### 3.3 Comparison of the principal curves method and scShaper

The principal curves algorithm is a widely used method in scRNA-seq trajectory inference and it is commonly applied at the final phase of trajectory inference to generate a continuous pseudotime. Since the performance differences observed in the benchmarking could be partially attributable to the different preprocessing steps, we investigated how the performances of principal curve method and scShaper compare when using the same preprocessing steps (**Section 2.2**). In this comparison, we used the implementation of the principal curves algorithm from the princurve R package, which is used by several popular trajectory inference methods, such as Slingshot and SCORPIUS. We performed the same benchmarking as in **Section 3.2** and the results suggested (**Fig. 4a**) that scShaper was better in terms of the overall score, as well as in terms of the accuracies of the cell ordering and the differentially expressed features (*p* < 0.05).

**Figure 4.**
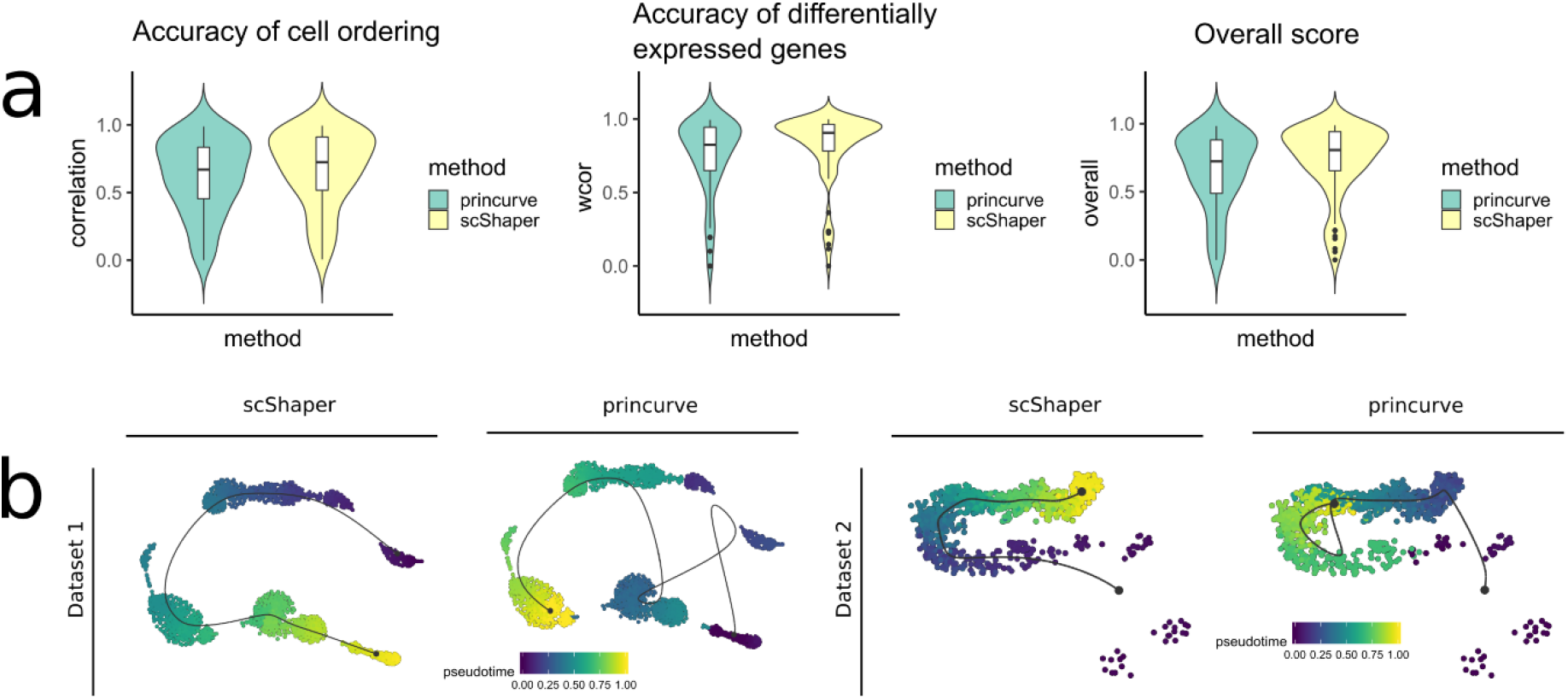
Comparison of the principal curve method (princurve) and scShaper for linear trajectory inference of scRNA-seq data. **(a)** The same benchmarking as in Section 3.2. **(b)** Trajectory visualizations of two simulated datasets, prosstt_linear_3 and prosstt_linear_6, from the comparison study. The coloring in each plot denotes the estimated pseudotime by the method. In all visualizations the trajectory line estimated using the dynplot R package is an approximation of the estimated pseudotime, and therefore the trajectory line bypasses some of the small subpopulations with fewer cells. In all the analyses we used the same dimensionality reduction workflow (**Section 2.2**) with both methods.

Next, we considered two datasets from the earlier benchmarking (**Fig. 3**) that were particularly challenging (prosstt_linear_3 and prosstt_linear_6) and visualized the estimated trajectories using the dynplot R package (**Fig. 4b**). For both datasets scShaper was able to generate trajectories that had almost perfect cell ordering accuracies (**Fig. 3d**), whereas the trajectories predicted by the principal curve method were highly inaccurate.

Finally, we investigated how accurately the two methods were able to predict the path through the middle of each of the simulated trigonometric trajectories, which included 2D and 3D spirals and the sine wave (**Section 2.4**). While both methods were able to accurately predict the path through the sine wave, only scShaper predicted accurately the ordering of the spiral trajectories (**Supplementary Fig. 5**).

### 3.4 Run time

The run time of scShaper mostly depends on three factors. First, it depends on the range of the *k* values used to estimate the discrete pseudotime. Our analysis shows (**Supplementary Fig. 1**) that the fast Rcpp implementation of the greedy algorithm in the scShaper R package can permute 99 distance matrices with *k* values from 2 to 100 in roughly 2.5 seconds. When the upper limit of the *k* values was increased to 200, the average run time was almost one minute. Second, the run time depends on the size of the dataset, i.e. how many features and samples it contains. **Figure 5** visualizes the results of a run time comparison for the five benchmarked methods, where the dyntoy tool was used to simulate 1000, 5000 and 10,000 cells with 1000 features. The run times of the five methods were on a largely similar scale, being at most a few minutes. For scShaper the analysis took 19 s and 120 s with 1000 and 5000 cells, respectively. It should be noted, however, that many of the methods had a relatively high memory requirement, and comp1 and scShaper were the only methods that successfully completed the analysis without memory failure with 10,000 cells on a laptop with 8 GB of RAM. For the largest dataset with 10,000 cells, the run time of scShaper was about four minutes. Finally, the run time also depends on the efficiency of the dimensionality reduction. Dimensionality reduction using PCA and *t*-SNE was clearly the slowest step of the scShaper’s workflow with all three datasets.

**Figure 5.**
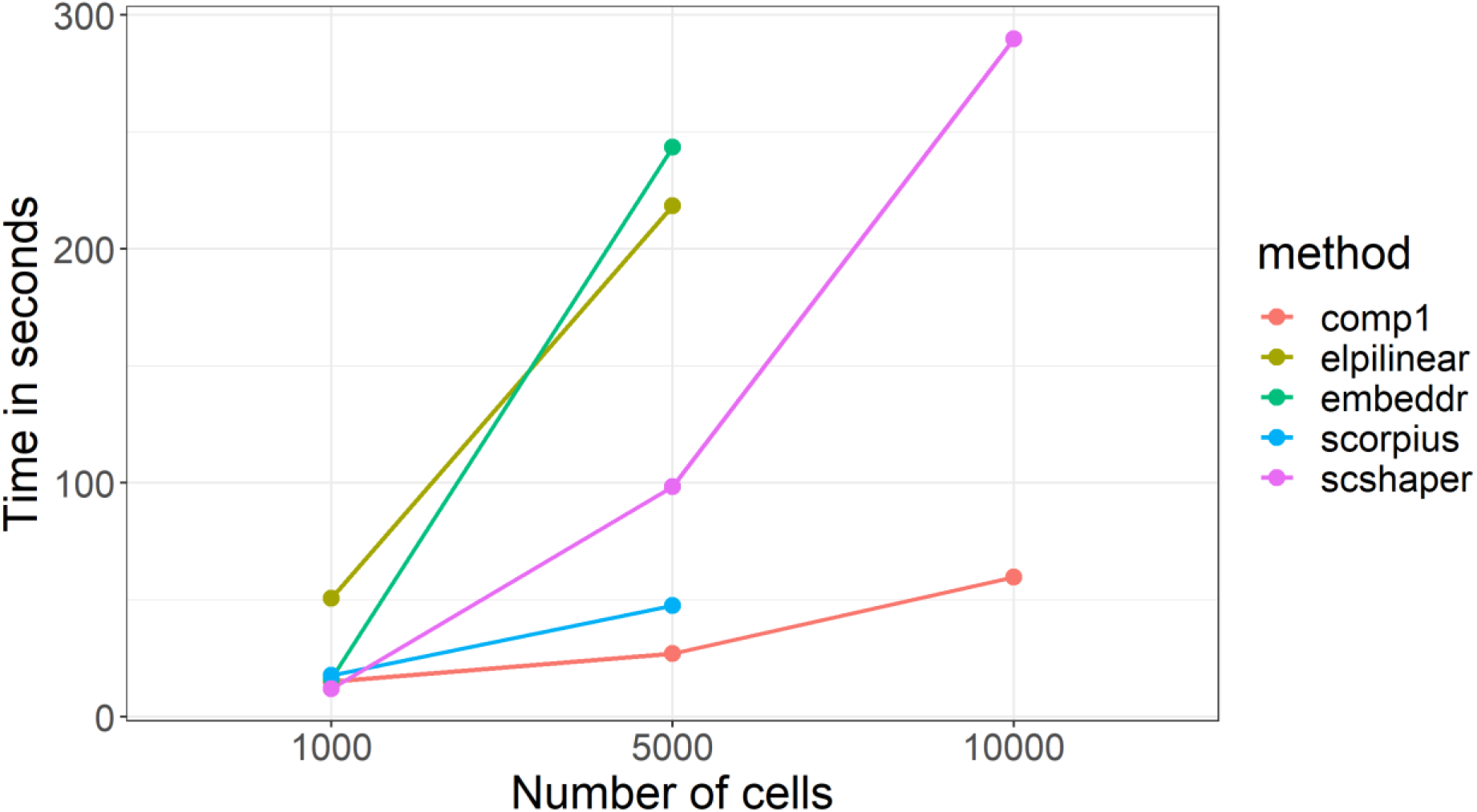
Run time comparison for the five benchmarked methods. With 10,000 cells, comp1 and scShaper were the only methods that were able to complete the analysis without memory failure on a laptop with 8GB of RAM.

## 4 Discussion

The current state-of-the-art trajectory inference methods, such as SCORPIUS and Slingshot, perform a single clustering, which is then used as the basis for inferring the topology of the trajectory using greedy graph theory algorithms, such as Kruskal’s algorithm. However, there are several issues with this approach that can deteriorate the performance. Greedy algorithms can often find only a local optimum, which is not necessarily the most optimal solution that accurately models the path through the real process. The clustering algorithms themselves, such as k-means and Gaussian mixture model (GMM), are stochastic, and the estimation of the number of clusters is known to be very challenging. All these issues make trajectory modelling a challenging task, which can be sensitive to small fluctuations in the input data and the model parameters. In addition, the principle curves algorithm, which is commonly used in the final step to smoothen each lineage of the trajectory, can significantly alter the path given as prior information and oversimplify the trajectory by bypassing many important milestones.

In this paper, we introduced scShaper, a novel linear trajectory inference method that utilizes an ensemble learning approach that combines multiple discrete pseudotimes derived from multiple clusterings. Ensemble methods are machine learning methods that combine multiple solutions in order to build a model that performs better than the individual models. They have already been widely applied in scRNA-seq data analysis, especially in consensus clustering and cell type classification (Smolander et al., 2020; Lieberman et al., 2018). However, so far there have been few attempts to apply the principle of ensemble learning to trajectory inference, although the same challenges are present there as well.

The ensemble approach of scShaper combines multiple discrete pseudotimes derived from multiple clusterings using different numbers of clusters to build a more accurate trajectory. Instead of specifying a single number of clusters, the user only needs to select a range of cluster numbers (default from 2 to 100). For each clustering, we estimate a discrete pseudotime using a greedy algorithm that is a special case of Kruskal’s algorithm for finding MSTs. scShaper applies PCA to analyse linear dependencies between the discrete pseudotimes, and an ensemble solution is built based on the largest subset of linearly correlating pseudotimes. Finally, we use LOESS to perform smoothing and generate the final continuous smooth pseudotime. scShaper mitigates the shortcomings of the current state-of-the-art methods, e.g. SCORPIUS, that use a single clustering as the basis for building the trajectory, making it less sensitive to the stochasticity of the clustering and the greediness of the graph theory algorithms that find the shortest path through the clustering.

We repeated the same comprehensive comparison of scRNA-seq trajectory inference methods that was recently published, which involved a wide array of different performance metrics and datasets of real and simulated origin (Saelens et al., 2019). The comparison showed that scShaper achieved superior accuracy for differentially expressed genes, while still maintaining accurate cell ordering. Indeed, the overall performance score of scShaper was best for most of the simulators and the real data. This suggests that scShaper is able to maintain a high cell ordering accuracy and model more accurately subpopulations that are bypassed by other methods.

Another significant part of this work included comparing the principal curves method and the pseudotemporal ordering algorithm of scShaper. Since the principal curves algorithm has become so widely utilized in scRNA-seq trajectory inference, we compared it with scShaper under the same preprocessing steps. The results showed that scShaper outperformed the principal curves algorithm in terms of all three metrics that were used in the evaluation. Moreover, scShaper achieved excellent accuracy for different nonlinear trajectories, including spiral and polynomial functions, which the principal curves method failed to accurately model. Since many of the benchmarked methods utilize the principal curves algorithm, this strongly suggests that the performance boost of scShaper is not attributable to the differences in the dimensionality reduction, but to the differences in the pseudotemporal ordering.

scShaper is a relatively simple algorithm with few hyperparameters. The most important parameter is the range of different *k* values, i.e. the number of clusters in *k*-means clustering, for which the discrete pseudotimes are estimated. Adjusting the range may be necessary, if the trajectories significantly exceed the complexity of the trajectories considered in this work. This can for example happen when we add considerably more rounds to the spiral trajectories (**Section 2.4**). The run time of scShaper largely depends on the range of *k* values (**Section 2.6**). For a set of clusterings, where *k* ranges from 2 to 100, the shortest path estimation lasted only two seconds, but when the upper limit was increased to 200, the analysis was noticeably longer (^~^ 1 minute).

To conclude, scShaper is a new fast and accurate method for linear trajectory inference from single-cell RNA-seq data. Our comparison showed that it outperformed current state-of-the-art methods (SCORPIUS, Embeddr) that utilize the principal curves method and also two other accurate methods (ElPiGraph Linear, Component 1). We observed a particularly large improvement in the accuracy of the differentially expressed genes, which we hypothesized to be related to the over-smoothing behaviour of the principal curves algorithm, making the principal curves algorithm to bypass some of the trajectory milestones. Further analysis suggested that the performance boost of scShaper is indeed not attributable to the dimensionality reduction, but the ensemble approach for generating the continuous pseudotime. scShaper can be downloaded as an user-friendly R package at https://github.com/elolab/scshaper.

## Acknowledgements

The authors thank the Elo lab for fruitful discussions and comments on the manuscript.

## Availability of data and materials

scShaper is available as an R package at https://github.com/elolab/scshaper. The benchmarking data are available at https://zenodo.org/record/1443566#.XiL7FyPRaUl and the related code at https://github.com/elolab/scshaper-benchmarking.

