## Supplementary Information for "scShaper: ensemble method for fast and accurate linear trajectory inference from single-cell RNA-seq data"

#### Supplementary Figure 1

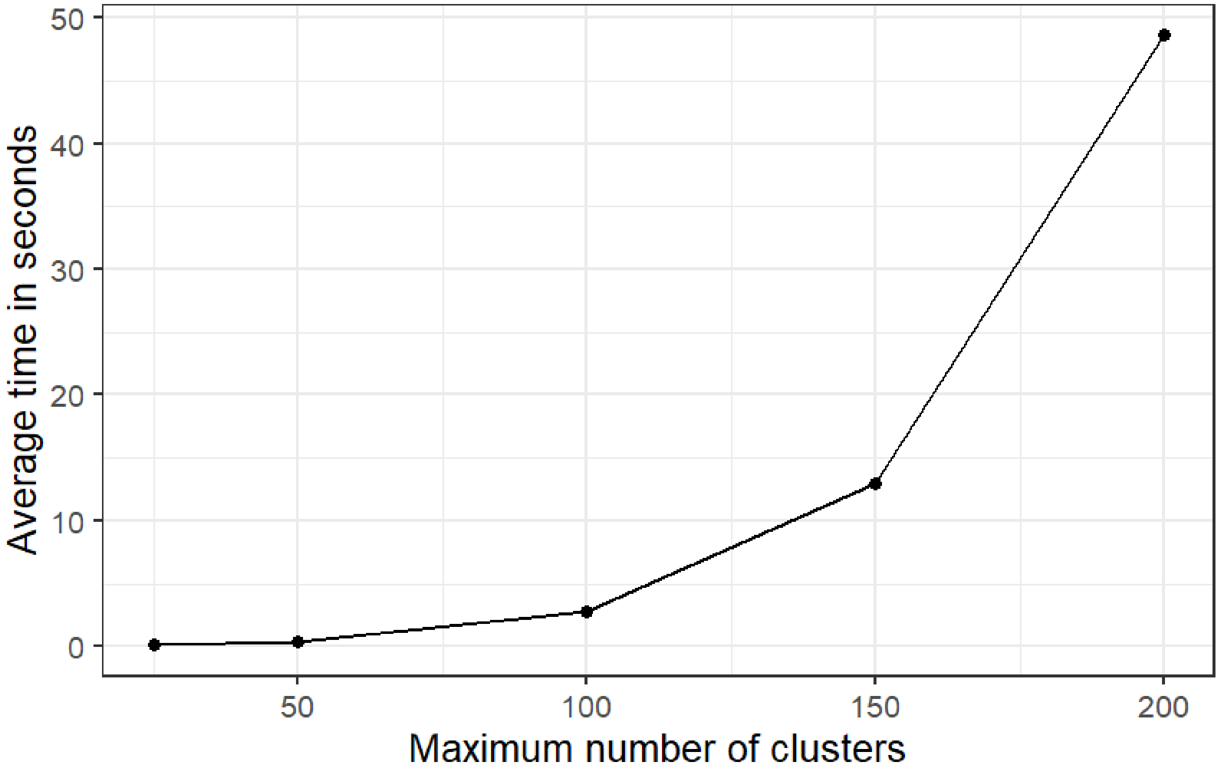


**Supplementary Figure 1.** Average run time in seconds of 100 repetitions of $k_{max} - 1$ greedy algorithm runs that solve the shortest Hamiltonian path problem for distance matrices with $k \in \{2, ..., k_{max}\}$ clusters, where $k_{max}$ is the maximum number of clusters. The greedy algorithm is implemented in Rcpp in the scShaper R package.

### Supplementary Figure 2

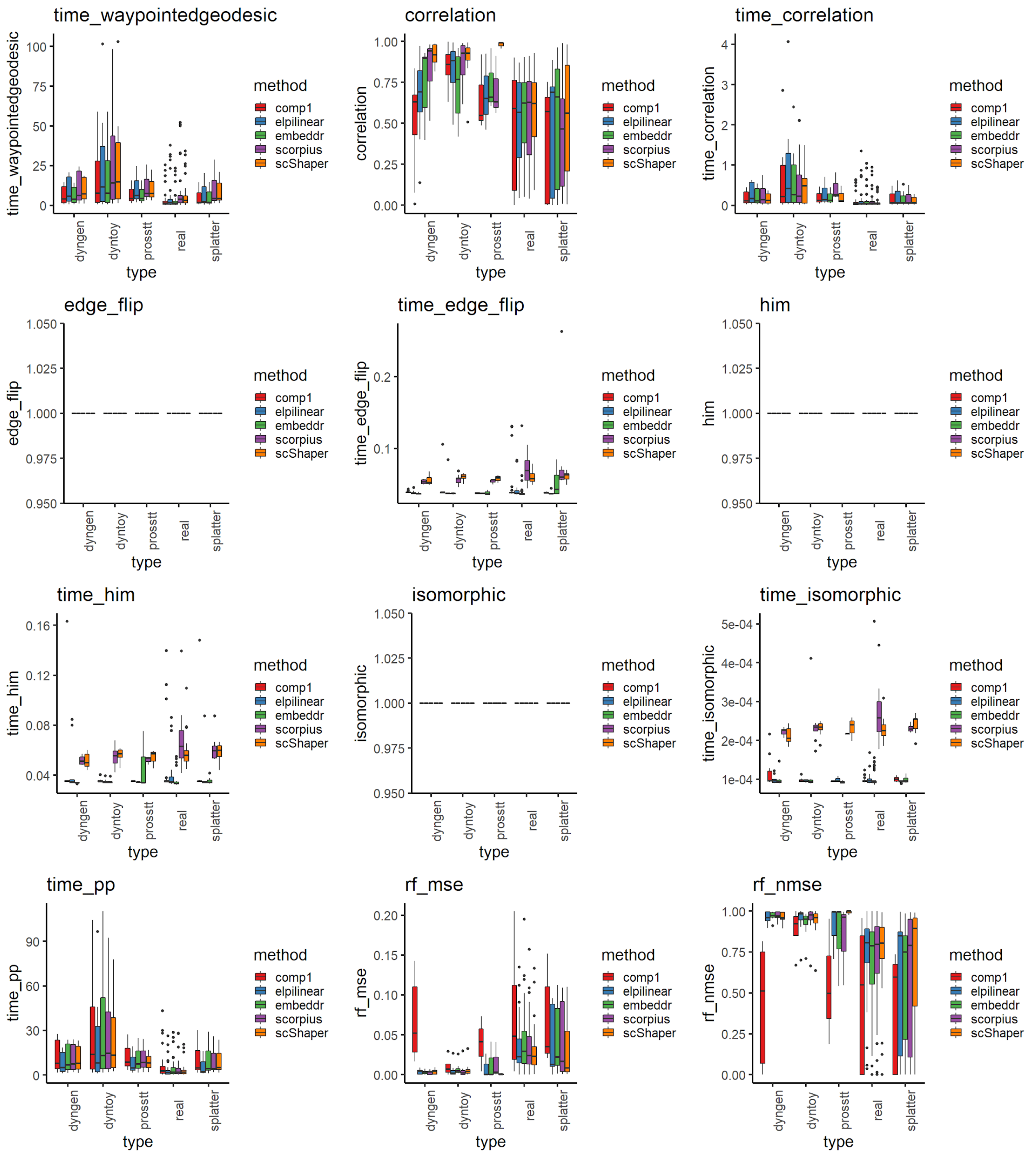


**Supplementary Figure 2.** The first twelve of the 31 benchmarking metrics computed using the dynbenchmark framework. See **Supplementary Figs. 3-4** for the visualizations of the rest of the metrics. With rf_mse (random forest mean squared error) a smaller value means better performance, whereas with the other measures a larger value means better performance. The metrics that measure topology and branching accuracy (HIM, isomorphic) are irrelevant in linear trajectory inference.

### Supplementary Figure 3

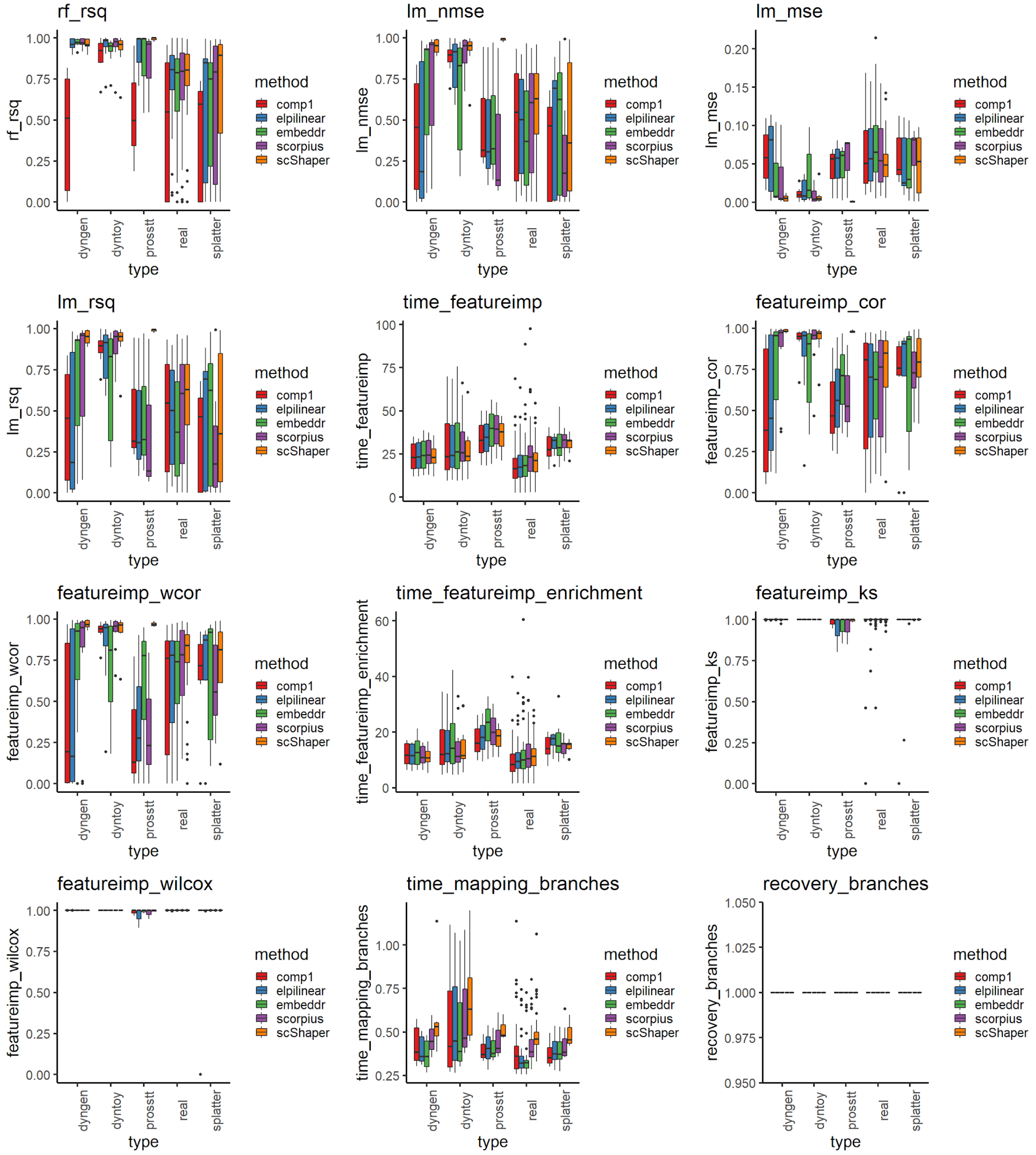


**Supplementary Figure 3.** The second twelve of the 31 benchmarking metrics computed using the dynbenchmark framework. See **Supplementary Figs. 2 and 4** for the visualizations of the rest of the metrics. With lm_mse (linear model mean squared error) a smaller value means better performance, whereas with the other measures a larger value means better performance. The metrics that measure topology and branching accuracy (recovery_branches) are irrelevant in linear trajectory inference.

### Supplementary Figure 4

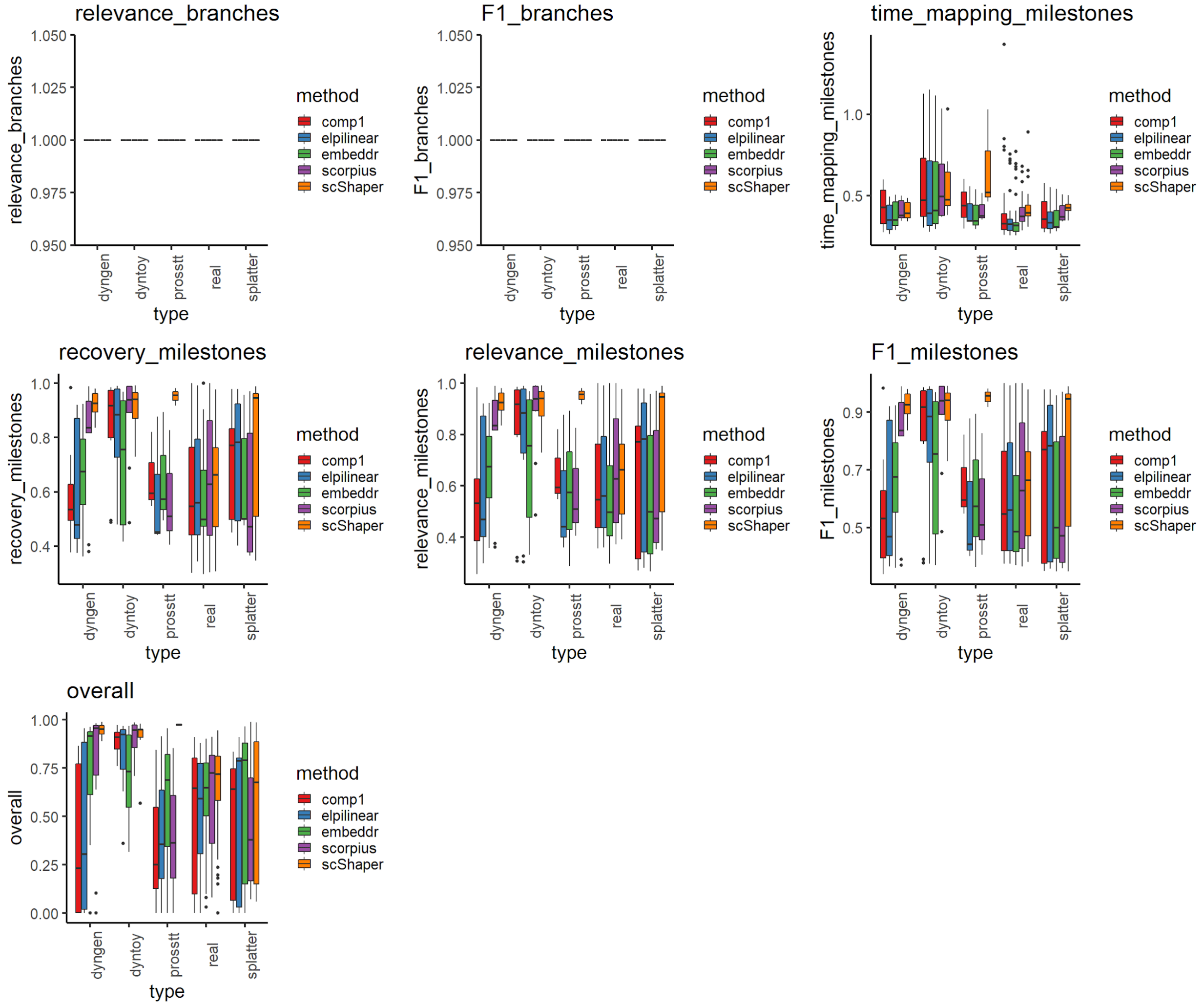


**Supplementary Figure 4.** The last seven of the 31 benchmarking metrics computed using the dynbenchmark framework. See **Supplementary Figs. 2-3** for the visualizations of the rest of the metrics. The metrics that measure topology and branching accuracy (relevance_branches, F1 branches) are irrelevant in linear trajectory inference.

### Supplementary Figure 5

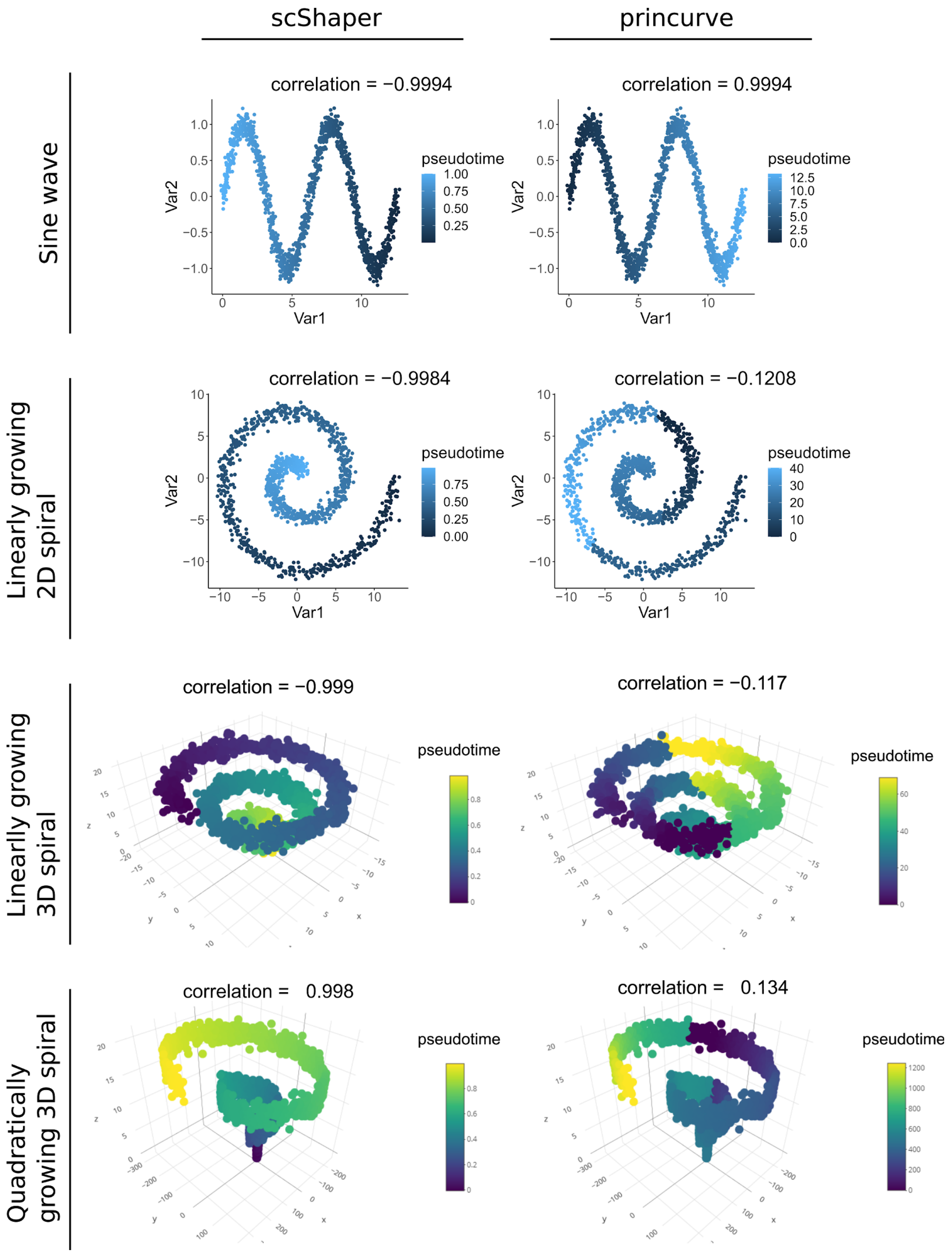


**Supplementary Figure 5.** Comparison of scShaper and the principal curve method from the princurve R package with data simulated using four trigonometric functions (**Section 2.4 of manuscript**). The correlation value above each plot denotes the Pearson correlation coefficient between the inferred and ground truth pseudotimes.
